# A scalable, reproducible and open-source pipeline for morphologically profiling image cytometry data

**DOI:** 10.1101/2022.10.24.512549

**Authors:** Maxim Lippeveld, Daniel Peralta, Andrew Filby, Yvan Saeys

## Abstract

Due to high resolution and throughput of modern image cytometry platforms, morphologically profiling generated datasets poses a significant computational challenge. Here, we present Scalable Cytometry Image Processing (SCIP), an image processing software aimed at running on distributed high performance computing infrastructure. SCIP is scalable, flexible, open-source and enables reproducible image processing. It performs projection, illumination correction, segmentation, background masking and extensive morphological profiling on various imaging types.

We showcase SCIP’s capabilities on three large-scale image cytometry datasets. First, we process an imaging flow cytometry (IFC) dataset of human white blood cells and show how the obtained features are used to classify cells into 8 cell types based on bright- and darkfield imagery. Secondly, we process an automated microscopy dataset of human white blood cells to divide them into cell types in an unsupervised manner. Finally, a high-content screening dataset of breast cancer cells is processed to predict the mechanism-of-action of a large set of compound treatments.

The software can be installed from the PyPi repository. Its source code is available at https://github.com/ScalableCytometryImageProcessing/SCIP under the GNU General Public License version 3. It has been tested on Unix operating systems. Issues with the software can be submitted at https://github.com/ScalableCytometryImageProcessing/SCIP/issues.

**Author Summary:** Cytometry is a field of biology that studies cells by measuring their characteristics. In image cytometry, this is done by acquiring images of cells. In order to gain biological insight from a set of images, an extensive amount of measurements are derived from them describing the cells they contain. These measurements include, for instance, a cell’s area, diameter, or the average brightness of the cell image. These measurements can then be analyzed using automated software tools to understand, for example, how cells respond to drug treatments, or how cells differ between a healthy and a diseased person. In this work, we present a novel software tool that is able to efficiently compute image measurements on large datasets of images. We do this by harnessing the power of high performance computing infrastructure. By enabling image cytometry researchers to make use of more computational power, they can more efficiently process complex and large datasets, paving the way to novel, fascinating biological discoveries.

## 2 Introduction

High-throughput imaging technologies collect detailed information on large volumes of cells enabling unprecedented insight into biological systems [20, 30, 16, 33]. Over the last few decades, advances in machine learning have greatly driven forward the potential of data generated by various bioimaging modalities. For example, cellular morphological profiles extracted from automated microscopy data have been combined with deep neural networks to repurpose high-throughput image assays for more efficient drug discovery[21]. Other machine learning methods have enabled stain-free classification of leukocytes in human whole blood samples using IFC data[26, 23], or have improved breast cancer detection using ultrasound imaging[25].

Applying machine learning methods on these datasets requires them to be digested into numerical profiles that describe the cell’s phenotype. These image-derived profiles contain hundreds of parameters describing cells in an unbiased way, enabling unexpected novel biological insights [30]. However, to keep on top of profiling the abundance of generated data, software solutions have to evolve together with imaging hardware. While in the past imaging data could be analyzed on a desktop computer, software running on a powerful server or high performance computing cluster is required to meet current and future demands.

Existing image cytometry analysis tools such as CellProfiler[18, 2], ImageJ[17, 9], QuPath[15], MCMICRO[34], and Orbit[29] are heavily used by researchers to obtain novel insights into complex imaging data. These tools perform, among other tasks, image segmentation and masking, and feature profiling. Table 1 gives an overview of how they compare to each other.

**Table 1:**
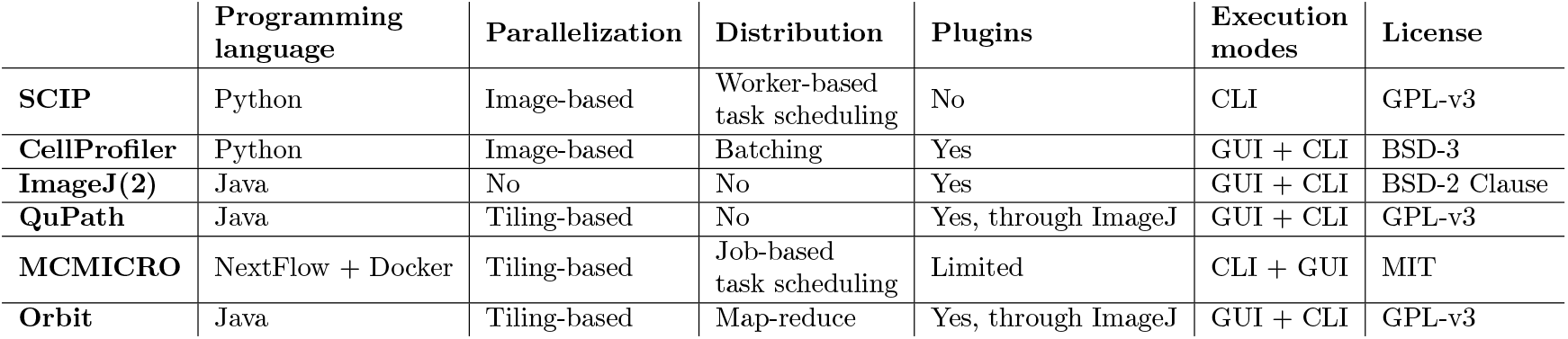
Overview of existing bioimage analysis tools and comparison to SCIP.

|  | Programming language | Parallelization | Distribution | Plugins | Execution modes | License |
| --- | --- | --- | --- | --- | --- | --- |
| **SCIP** | Python | Image-based | Worker-based task scheduling | No | CLI | GPL-v3 |
| **CellProfiler** | Python | Image-based | Batching | Yes | GUI + CLI | BSD-3 |
| **ImageJ(2)** | Java | No | No | Yes | GUI + CLI | BSD-2 Clause |
| **QuPath** | Java | Tiling-based | No | Yes, through ImageJ | GUI + CLI | GPL-v3 |
| **MCMICRO** | NextFlow + Docker | Tiling-based | Job-based task scheduling | Limited | CLI + GUI | MIT |
| **Orbit** | Java | Tiling-based | Map-reduce | Yes, through ImageJ | GUI + CLI | GPL-v3 |

While these tools have extensive functionality, they have, except for Orbit, a limited focus on scalability and focus more on usage with a graphical user interface on local workstations. This makes them easily adoptable by a diverse audience, but introduces challenges when we need to scale to large datasets with millions of cells.

In order to be properly scalable, work needs to be distributed over multiple, interconnected machines. For example, in CellProfiler and MCMICRO, scaling is achieved by statically partitioning the dataset and processing the partitions in parallel on one or more machines[24]. However, this *split-apply-combine* approach is less flexible, since no communication is possible between processes executing the pipeline. For instance in CellProfiler, it is not possible to run different parts of a pipeline on dedicated hardware, such as running segmentation on a GPU-accelerated machine.

Orbit does provide distributed processing by implementing a tile-based MapReduce framework based on Apache Spark. However, Orbit is implemented in Java, which makes it less suited for fast adaptation of state-of-the-art techniques that are often developed in Python. Addition-ally, just like QuPath, which is also implemented in Java, it can not benefit from the rich data science ecosystem in Python.

In this work, we present Scalable Cytometry Image Processing (SCIP), a software package implemented in Python for morphologically profiling large-scale image cytometry datasets. SCIP is scalable, flexible, open-source and enables reproducible image processing.

By building SCIP on top of Dask [11], a framework for distributed computing in Python, it scales from laptops to high performance computing clusters, enabling profiling of small to large-scale datasets with millions of cells.

Because of SCIP’s design and implementation, it is more flexible than existing tools. First, by combining Dask’s smart task scheduling and SCIP’s modularity it is possible to execute parts of the pipeline on specialized hardware such as a GPU. Secondly, SCIP comes with all of Dask’s benefits such as fault tolerance, load balancing across workers and access to distributed file systems. Finally, Dask clusters can be set up in a large variety of compute environments, and can be dynamically scaled up or down with availability of resources.

SCIP also enables reproducible image processing. First, configuration of the software happens through easily shareable YAML configuration files, making it easy to rerun SCIP with the same settings. Secondly, its source is available under the GPL-v3 license. All code is available on Github at https://github.com/ScalableCytometryImageProcessing/SCIP. Finally, the software is easily installable from the PyPi (https://pypi.org/project/scip/) repository.

To showcase SCIP’s flexibility, we apply it on large-scale image cytometry datasets in three use cases. The first and second datasets characterize human whole blood cells using an IFC and an automated microscopy platform, respectively. The third is BBBC021, a publicly available high-content screening dataset characterizing the response of breast cancer cells to various small molecule treatments.

## 3 Design and implementation

In this section, we give an overview of the image profiling pipeline implemented in SCIP, discuss how it can be efficiently parallelized with graph-based task scheduling, and show how SCIP is implemented using the Dask framework.

### 3.1 SCIP implements an extensive image profiling pipeline

To profile a dataset, SCIP implements a pipeline consisting of a projection, illumination correction, segmentation, masking and profiling step. In the following, we discuss for each step what is computed and how it is concretely implemented.

The input is a matrix of pixels *D* which encode the measured signal intensity. All datasets have dimensions *X, Y*, channel (*C*) and index (*I*). In multi-focal datasets, a fifth *Z* dimension indexes the different focal planes. We refer to each position *i* on the index as image (*d*_*i*_) *∈* ℝ^*c×z×x×y*^. Each image *d*_*i*_ also has meta data associated with it, which are a number of categorical or ordinal values.

#### Projection

For a multi-focal dataset, the images are projected onto one plane as shown in Equation 1.

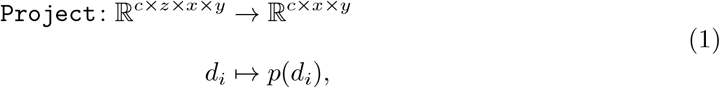

where *p* is a projection function over dimension *z*, such as the max operator that outputs the maximum value over the focal planes on each (*c, x, y*) location in *d*_*i*_. SCIP provides implementations for min, mean, max and median projection.

#### Illumination correction

Next, illumination correction transforms images to mitigate the influence of non-homogeneous illumination. Here, we focus on retrospective correction, which combines data from all images acquired in a batch to produce an illumination correction function (ICF). The function is applied to each image to produce a corrected version (Equation 2)[8].

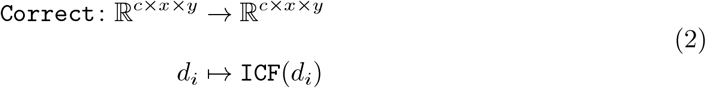

SCIP provides the ICF proposed by Jones et al. This approach has also been used to process the dataset in the use case of Section 4.1.2. In this implementation, the ICF is computed by producing a smoothed average of all images per channel per experimental batch. For a dataset *D* with *m* experimental batches *D*_*j*_ the ICF is computed for each batch as follows

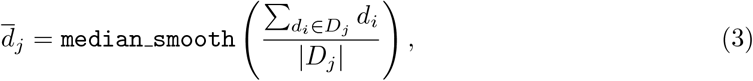

where `median_smooth` is a median filter. The ICF is applied by dividing each image by the corresponding 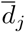.

#### Segmentation

In datasets where each field of view contains many cells, instance segmentation is required to identify regions containing single cells. The segmentation operation is defined in Equation 4.

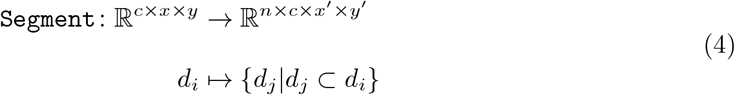

In SCIP, Cellpose is used for segmentation. It is a deep learning-based generalist algorithm for cellular segmentation of microscopy data[32], which can be applied on a wide variety of input images. It achieves state-of-the-art segmentation results by implementing a novel model design for cell segmentation based on the principles of watershed segmentation, as well as a new training dataset containing a wide variety of images with manually segmented objects.

#### Masking

In IFC-like data, each *d*_*i*_ is expected to contain one cell. The masking operation (Equation 5) identifies the region containing the cell’s signal in each channel of the image.

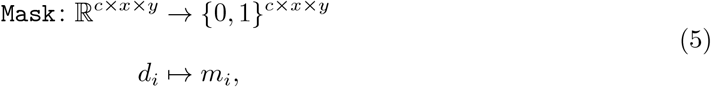

where *f* is a masking function that sets each (*x, y*) location to either 0 (background) or 1 (foreground) based on some decision function. SCIP has implementations for threshold and spot masks, which mask the entire cell area or the brightest spots in it, respectively.

#### Profiling

Finally, the images are profiled. This operation (Equation 6 and 7) maps the pixel and mask data of *d*_*i*_ onto a feature vector that describes characteristics such as shape, texture or signal intensity of the object captured in *d*_*i*_.

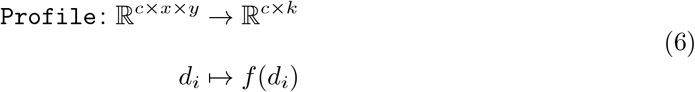

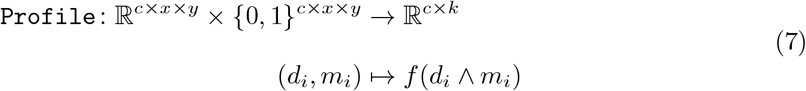

Three types of features are derived from the pixel and mask data: intensity, shape and texture. Table 2 gives an example of some of the computed features. Supplementary Section S1 describes the features in detail. All features are computed for every channel in the image using the channel-specific mask, as well as a union of all masks. The intensity and texture features are also computed on the pixels on the edge of the mask.

**Table 2:** Examples of features computed by SCIP. The last column indicates whether features are also computed on edge pixels of an object’s mask.

| Type | Examples | Computed on edge |
| --- | --- | --- |
| Intensity | minimum, maximum,<br>standard deviation,<br>skew, kurtosis | Yes |
| Shape | eccentricity, area,<br>diameter, solidity,<br>major axis length | No |
| Texture | Grey-level co-occurrence matrix<br>(contrast, correlation)<br>Sobel map<br>(max, standard deviation) | Yes |

### 3.2 Graph-based task scheduling enables fine-grained control over distributed pipeline execution

A naive parallel implementation of an image profiling pipeline processes all images *d*_*i*_ in dataset *D* using a number of isolated processes that execute the full function *f*, defined as follows

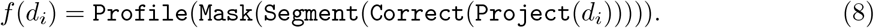

However, this approach is not flexible due to the lack of communication and cooperation between the processes.

A way to overcome these limitations, is by using graph-based task scheduling. In this approach, a directed acyclic graph (DAG), called the task graph, is constructed that encodes all tasks to be executed on the dataset as nodes, and dependencies between tasks as edges. A task graph executing the same functions as Equation 8 is shown in Figure 1.

**Figure 1:**
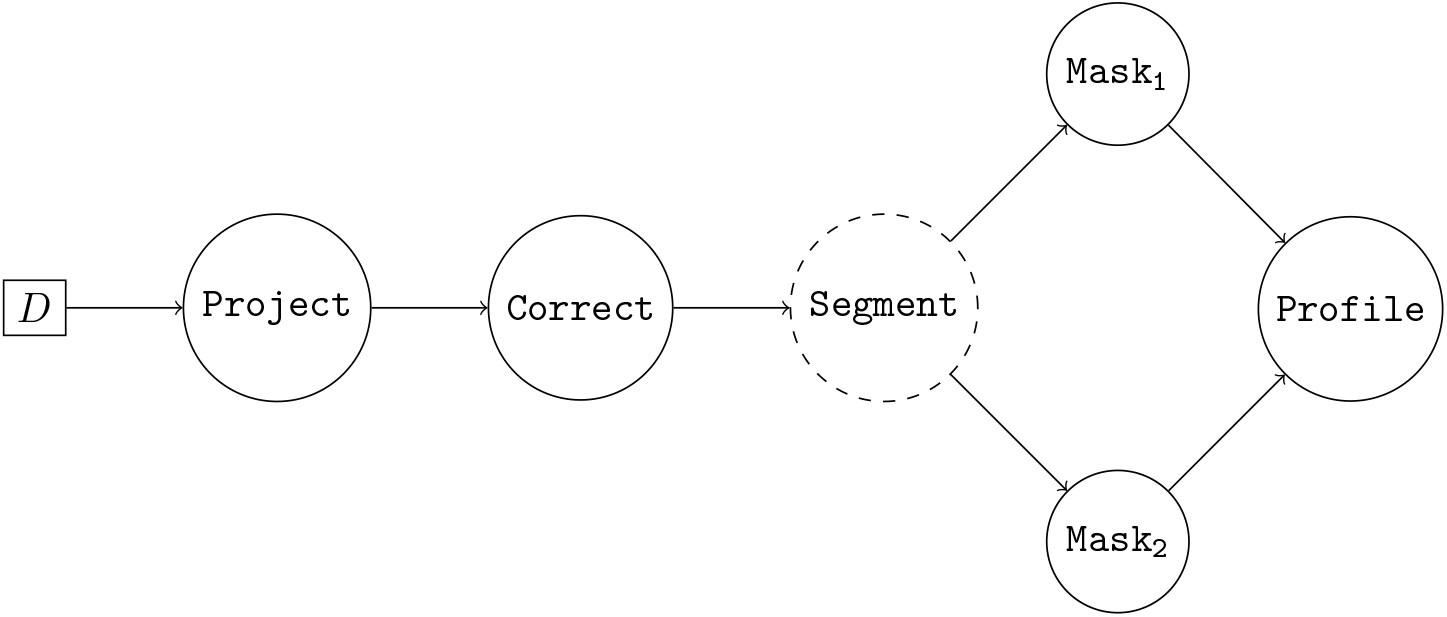
Basic task graph encoding the operations of Equation 8 to process dataset *D*. Nodes represent operations, edges represent dependencies between operations. Nodes with dashed borders have specific resource requirements.

The task graph is analyzed by a scheduler which instructs the worker processes to execute individual tasks, leveraging parallelization and reusing results where possible. The dynamic scheduler can also keep track of what each worker is executing so that it can resubmit tasks in the event of a worker fault, and so that it can rebalance the workload if necessary.

A number of frameworks implement graph-based task scheduling, including Apache Spark[13], Ray[19] and Dask. We chose to implement SCIP using the latter, because it is written in Python and it implements functionality that is a natural fit to the pipeline introduced above.

### 3.3 Dask-based implementation of SCIP enables scalability to large datasets

#### Algorithm 1: Extract image profiles from image dataset *D* with SCIP

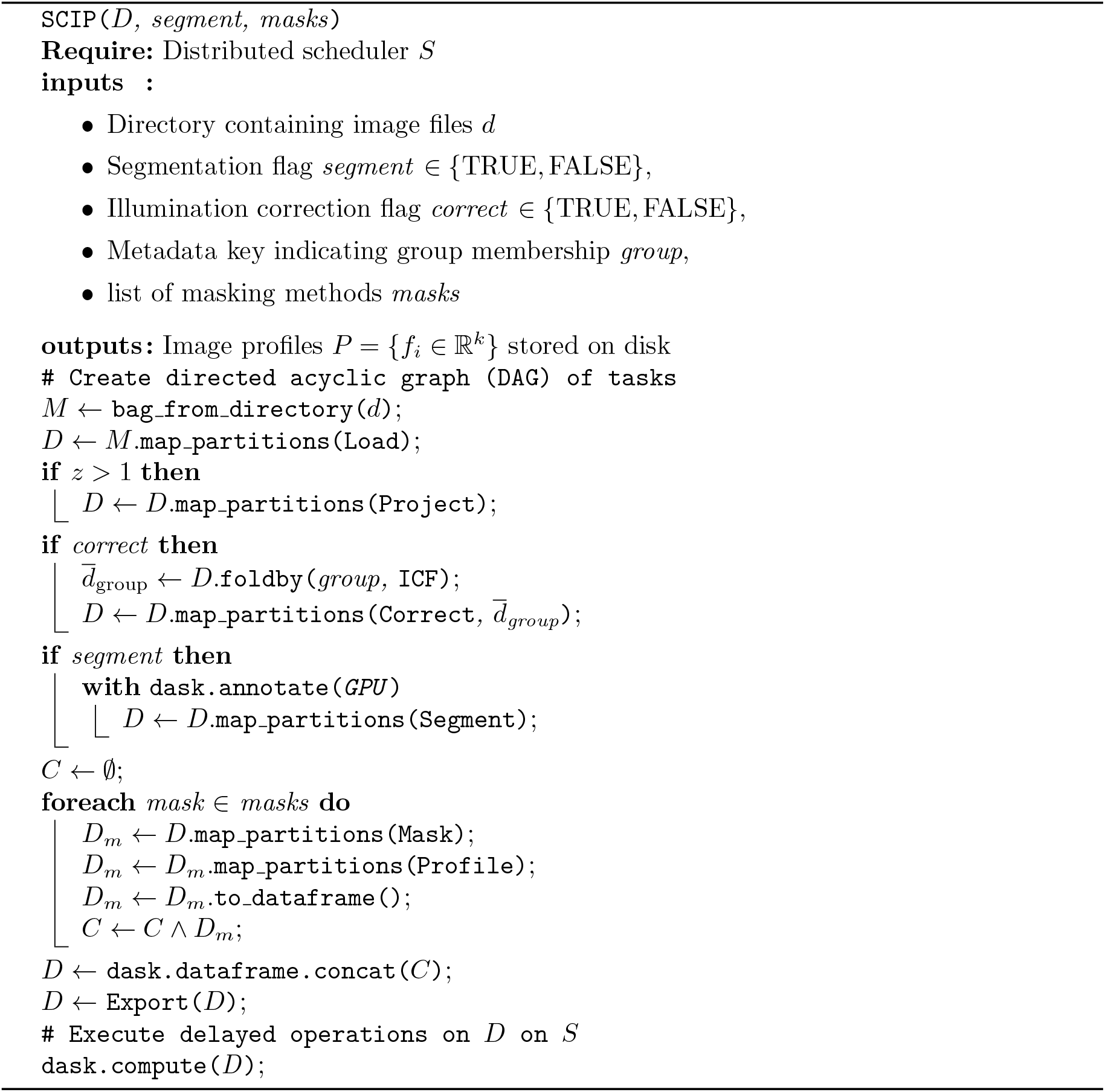

Algorithm 1 shows how the pipeline is implemented using operations on distributed Dask collections. The collections are used to construct a task graph that applies the functions defined in Section 3.1 on the input images. The task graph is sent to the Dask distributed scheduler which executes it as efficiently as possible on the available computational resources.

First, paths pointing to input files are stored in a Dask Bag *B* of dictionaries holding the meta data about each image *d*_*i*_. In the loading operation, pixel data is added to each dictionary in the Bag. By using a Bag of dictionaries, we can handle images with varying *x* and *y* dimensions and we can keep the meta data linked to the pixel data.

Illumination correction is available in SCIP through a distributed implementation of the method introduced in 3.1. It is implemented in three steps: First, a foldby operation is used to compute an average image per group. This is done by computing a distributed sum of all images and dividing the sum by the number of images in each group. Secondly, each averaged image is smoothed using a median filter. To reduce the memory footprint of the median filter, the image can be downscaled prior to filtering. Note that within each group, images need to have the same dimensions to apply correction.

Next, instance segmentation is performed using watershed or CellPose segmentation. All channels of the image are segmented separately; objects detected in different channels are assigned to the overlapping parent object detected in the user-defined parent channel. Finally, the segmentation step requires restructuring *B* from a *per-file* to a *per-object* Bag, that is, where *B* contains one element per file, bag *B*′ contains one element per cell.

Dask allows for tasks to be annotated with required resources, such as a GPU, which is specifically interesting for GPU-accelerated segmentation methods. The scheduler will only assign these task to a worker that has the requested resource. Figure 2 shows the stream of tasks executed by the workers, three of which have a GPU resource. It illustrates how heterogeneous computational resources can be used efficiently in SCIP.

**Figure 2:**
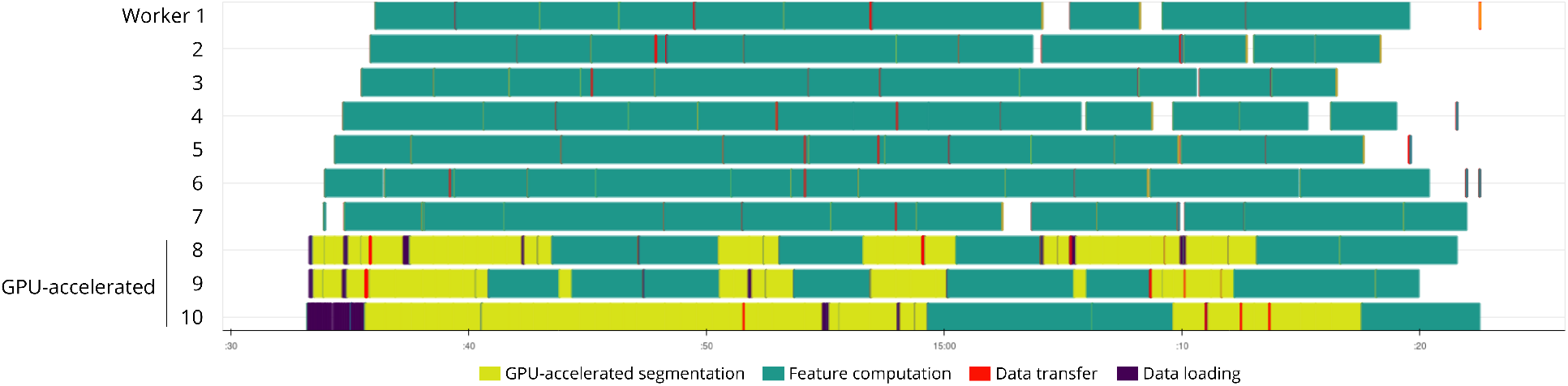
The Dask task stream of a microscopy dataset being processed with SCIP showing how heterogeneous computational resources can be used efficiently by distributing steps of the pipeline across the cluster. In this pipeline, segmentation is done with CellPose on three GPU-accelerated nodes.

Multiple masks can be computed to produce specific views on the images, such as a threshold and a spot mask. Figure 3 shows how this multi-mask approach translates to the task graph. Dask’s dynamic task scheduler will cache as much of the loaded images in distributed memory as possible to avoid re-loading images from disk for each mask.

**Figure 3:**
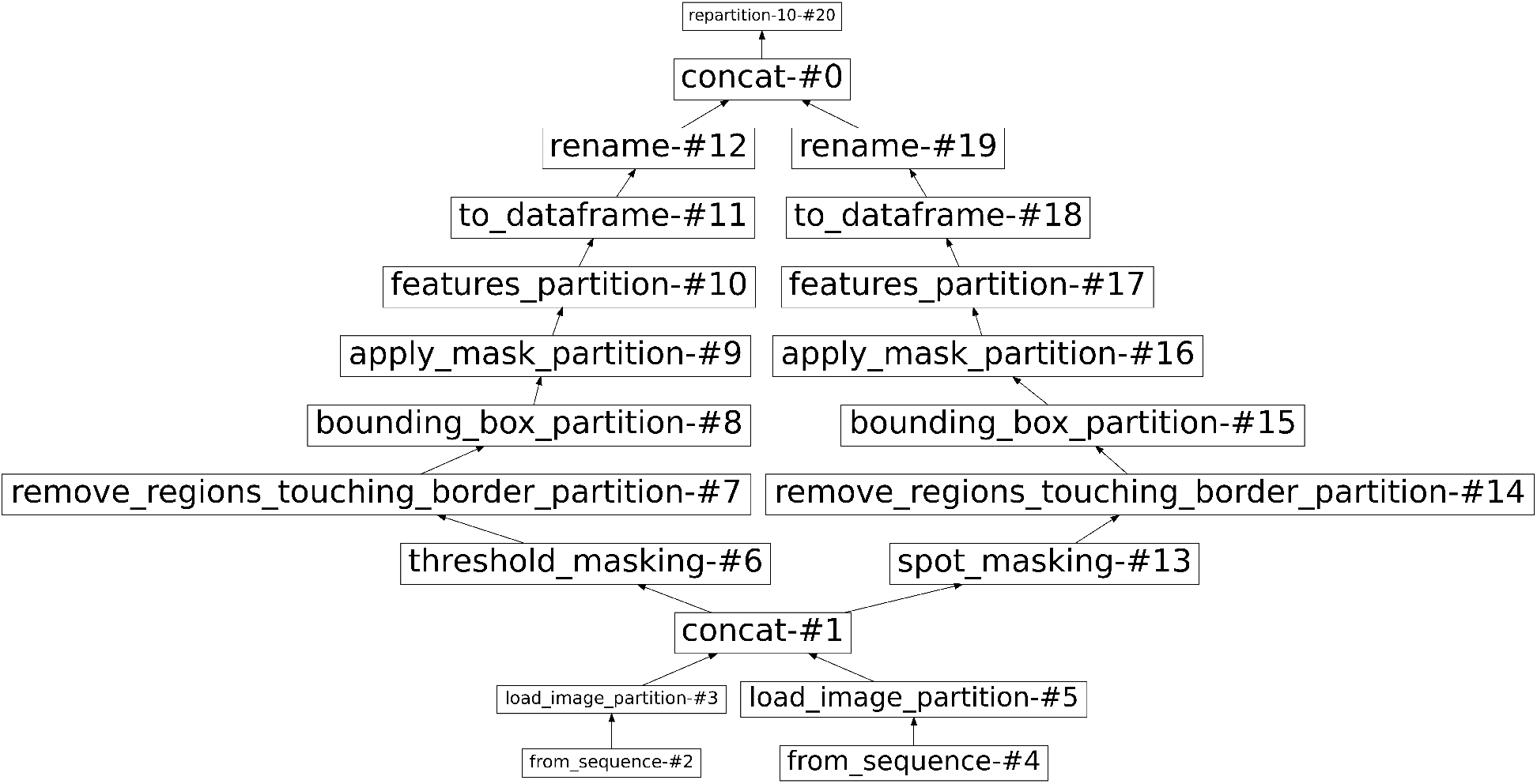
Dask task graph visualizing the operations to be executed on the input images. This particular graph loads in images from two files, computes two different masks for them (a threshold mask and a spot mask), and derives profiles from the images combined with the masks.

After segmentation and masking, the profiling operation is applied to *B*. This operation converts the Bag into a DataFrame containing a profile for each detected cell. The profiles are then exported to AnnData h5ad or to Parquet-files, to provide interoperability with the ScanPy[22] library and other programming languages, respectively. It is also possible to export segmentation results for inspection.

## 4 Results

In the following sections we show SCIP’s flexibility and scalability by applying it on three use cases, and on three performance experiments measuring runtime and memory usage in multiple configurations.

Experiments requiring GPU-acceleration were run on a machine with 4 8-core Intel Xeon Silver 4110 CPU’s and an NVIDIA GeForce RTX 3090 GPU. All other experiments were run on the Ghent University high performance computing Tier-2 doduo cluster. Each node in this cluster has 2 48-core AMD EPYC 7552 CPU’s and 250GiB of memory. Experiments were run on one to five nodes.

### 4.1 Use cases

We demonstrate the use of SCIP on three datasets. All necessary Jupyter Notebooks to reproduce our results are provided together with a Snakemake[31] workflow on Github at https://github.com/ScalableCytometryImageProcessing/SCIP-use-case-workflows. Datasets for use cases 1 and 3 were generated at Newcastle University. All datasets are publicly available for download; details are provided in their respective sections.

#### 4.1.1 Stain-free classification of human white blood cells with imaging flow cytometry

In [26] we introduced a human white blood cell dataset acquired on a Luminex ImageStream - MKII IFC platform, and compared stain-free classification performance of 8 white blood cell sub-types using various machine learning models. Models were trained on feature profiles extracted from stain-free images (two bright-field images, and one dark-field) using the proprietary IDEAS software supplied by Luminex. We concluded that a gradient boosting ensemble[10] performed best. Here, we redo the analysis with SCIP using the original dataset and on an extended version that has one extra sample.

We show with this use case that SCIP quickly and efficiently profiles IFC data without the proprietary IDEAS software or the suboptimal IFC solution proposed by CellProfiler. In particular, the latter requires a tiling step, which introduces limitations in the way the data must be handled. By removing the necessity for tiling, SCIP is less error-prone and has fewer parameters.

The dataset consists of 3 human whole blood samples, each stained with 9 fluorescent markers. Each acquired image has 12-channels; 9 fluorescent, 2 brightfield and 1 darkfield channel. A ground truth label was assigned to all cells with manual gating on the fluorescent stains. We refer to [26] for details on the data acquisition and gating procedure. All data is accessible under study S-BIAD452 on the Bioimage Archive at https://www.ebi.ac.uk/biostudies/studies/S-BIAD452.

To profile the dataset with SCIP, we exported 16-bit tiff images from the IDEAS software and stored them in the Zarr format [27]. Feature profiles were extracted for 247 993 12-channel images masked with both a spot and threshold mask (see Figure 4). Profiling took 2 hours and 29 minutes using 16 workers, 4338 features were computed in total per image. After filtering out doublets and debris based on the derived profiles, 233 262 cells remained for classification.

**Figure 4:**
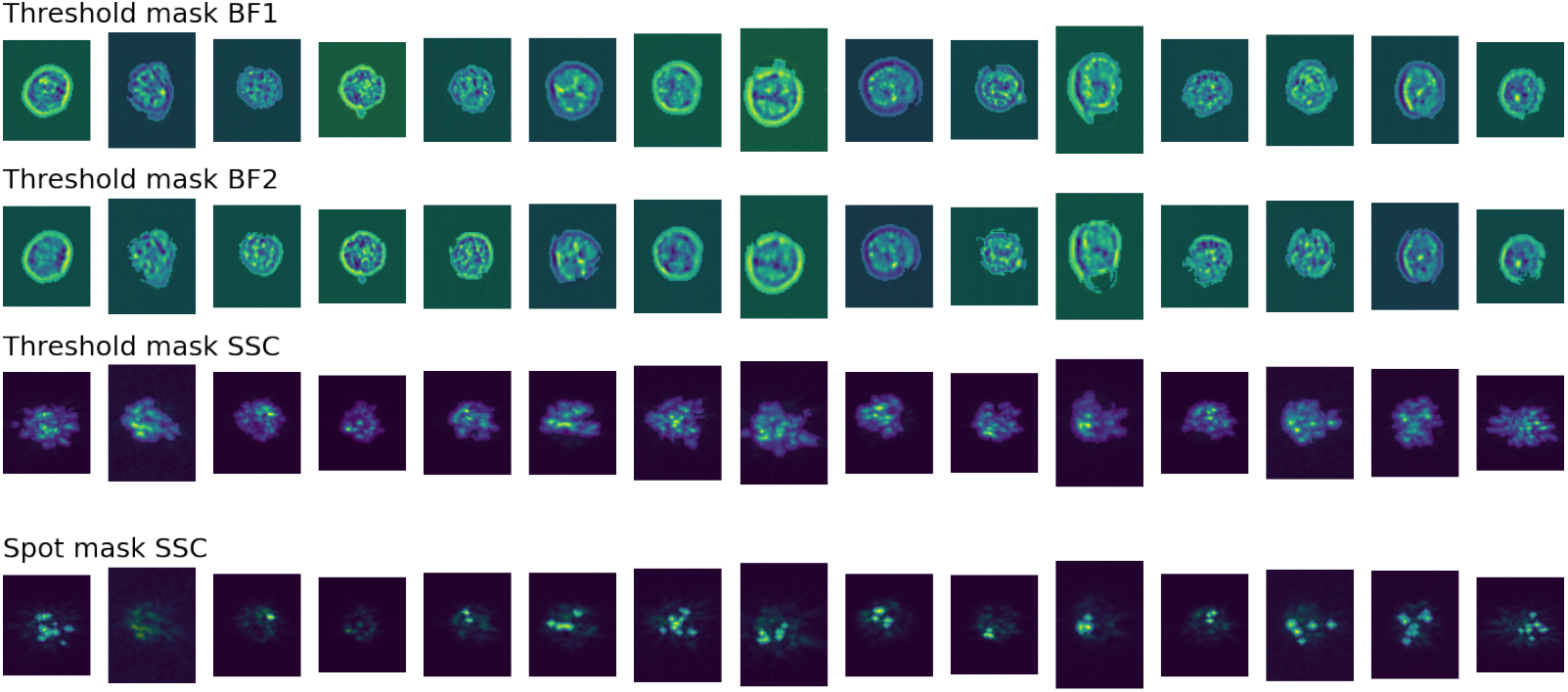
Example images of threshold and spot masks computed by SCIP for brightfield 1 (BF1), brightfield 2 (BF2) and darkfield (SSC) channels.

Using SCIP features we achieved a cross-validated balanced accuracy of 0.827 with standard error 0.002, compared to 0.853 (standard error 0.002) with IDEAS features (see Table 3) on the original dataset. The confusion matrices in Figures 5a and 5b show a similar pattern as in with major white blood cell type groups (lymphocytes, monocytes, neutrophils, eosinophils) being correctly distinguished, but subtypes, such as CD4+ versus CD8+ T-cells, confusing the model. We also present classification results on the extended dataset in Supplementary Section S2.1.2. Methodological details on the classification can also be found in Supplementary Section S2.1.2.

**Table 3:** Classification metrics for cross-validated stain-free leukocyte classification with an eXtreme Gradient Boosting classifier using features derived from SCIP and IDEAS.

| Metric Software | balanced accuracy |  | f1 macro |  | precision macro |  |
| --- | --- | --- | --- | --- | --- | --- |
|  | IDEAS | SCIP | IDEAS | SCIP | IDEAS | SCIP |
| Test | 0.853 (0.002) | 0.824 (0.002) | 0.800 (0.002) | 0.767 (0.007) | 0.770 (0.003) | 0.733 (0.008) |
| Train | 0.917 (0.004) | 0.921 (0.013) | 0.852 (0.004) | 0.849 (0.019) | 0.816 (0.004) | 0.807 (0.019) |

**Figure 5:**
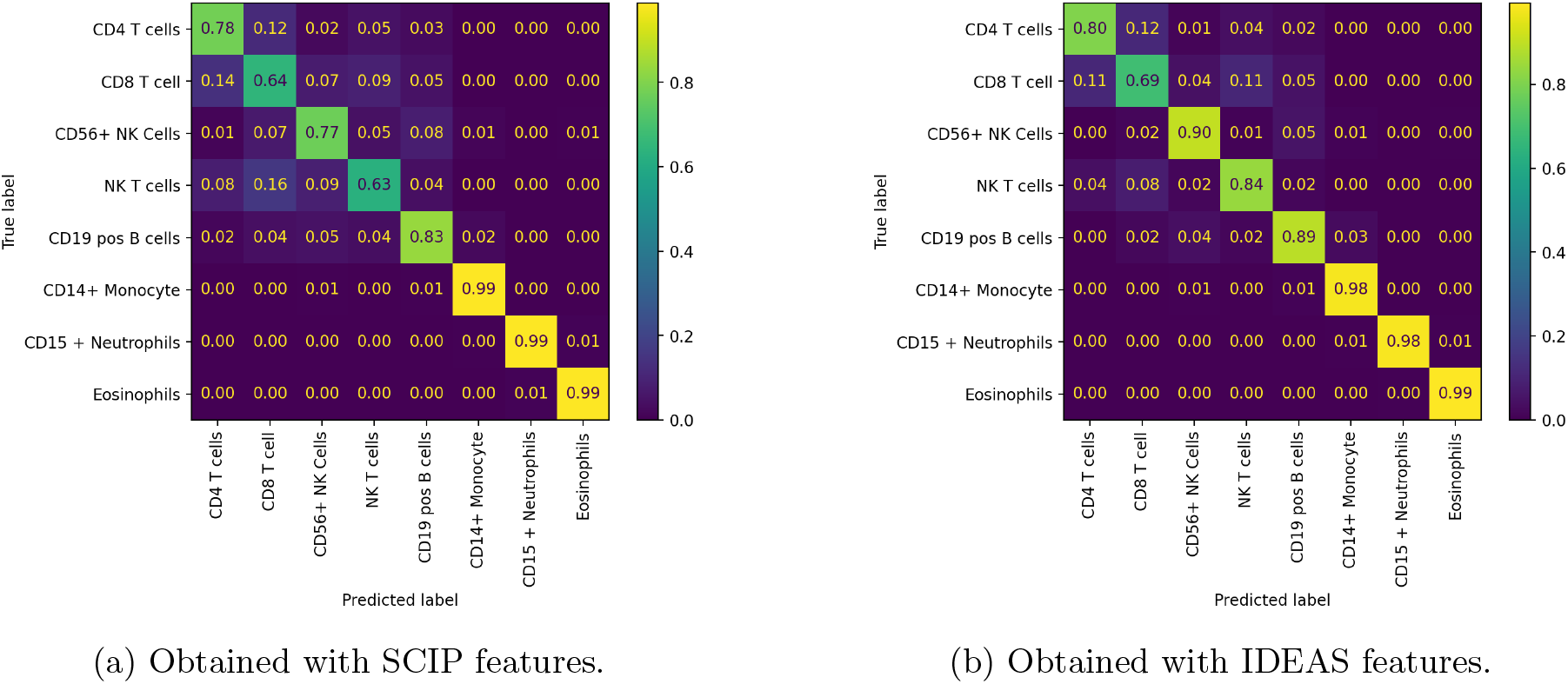
Classification results for cross-validated stain-free leukocyte classification with an eXtreme Gradient Boosting classifier.

We acknowledge that with IDEAS features we obtain slightly better classification results, however, it is not our goal to outperform IDEAS. We rather want to show that we can obtain similar performance using our scalable and open-source software.

#### 4.1.2 Predicting mechanism of action for various small-molecule treatments of MCF-7 breast cancer cells

We use SCIP to process data from a high-content screening experiment in which cells from a breast-cancer model system (MCF-7) were treated with a mechanically distinct set of targeted and cancer-relevant cytotoxic compounds. The dataset provides a basis for testing image-based profiling methods with respect to their ability to predict the mechanisms of action (MOA) of a treatment. The image dataset BBBC021v1[4] is available from the Broad Bioimage Benchmark Collection[5].

We show in this use case that it is possible to perform MOA prediction to a high degree of accuracy using morphological features derived by SCIP. Thanks to SCIP’s scalable design this can be done efficiently even for the large-scale dataset at hand, and when using GPU-accelerated segmentation algorithms, or algorithms which require aggregations over the dataset, such as illumination correction.

The dataset contains cells treated for 24 hours with 113 compounds at 8 concentrations. The cells were fixed, stained for DNA, F-actin, and *β*-tubulin, and imaged by fluorescent microscopy. In total, there are 39,600 image files (13,200 fields of view imaged in three channels). After acquisition a subset of the compound-concentrations were categorised into 12 primary MOAs. 6 of the 12 were identified visually, the remaining were defined based on the literature.

Several classification results on the MOA prediction task have been published, which all follow similar steps: (i) profile the cells using CellProfiler, (ii) summarize the profiles per treatment, and (iii) use a nearest-neighbor classifier to predict the MOA for a treatment. Singh et al. (2014) also proposed to perform illumination correction prior to profiling.

To obtain classification results we profiled the dataset using SCIP with illumination correction, and Cellpose segmentation. It took 17.6 hours to extract profiles for 2.18 million cells using 11 workers of which 4 were GPU-accelerated for segmentation. 470 127 cells remained when keeping only cells from the subset of images annotated with a MOA. The CellProfiler pipeline identified 454 793 cells. Table 4 shows results of existing approaches to MOA prediction.

**Table 4:**
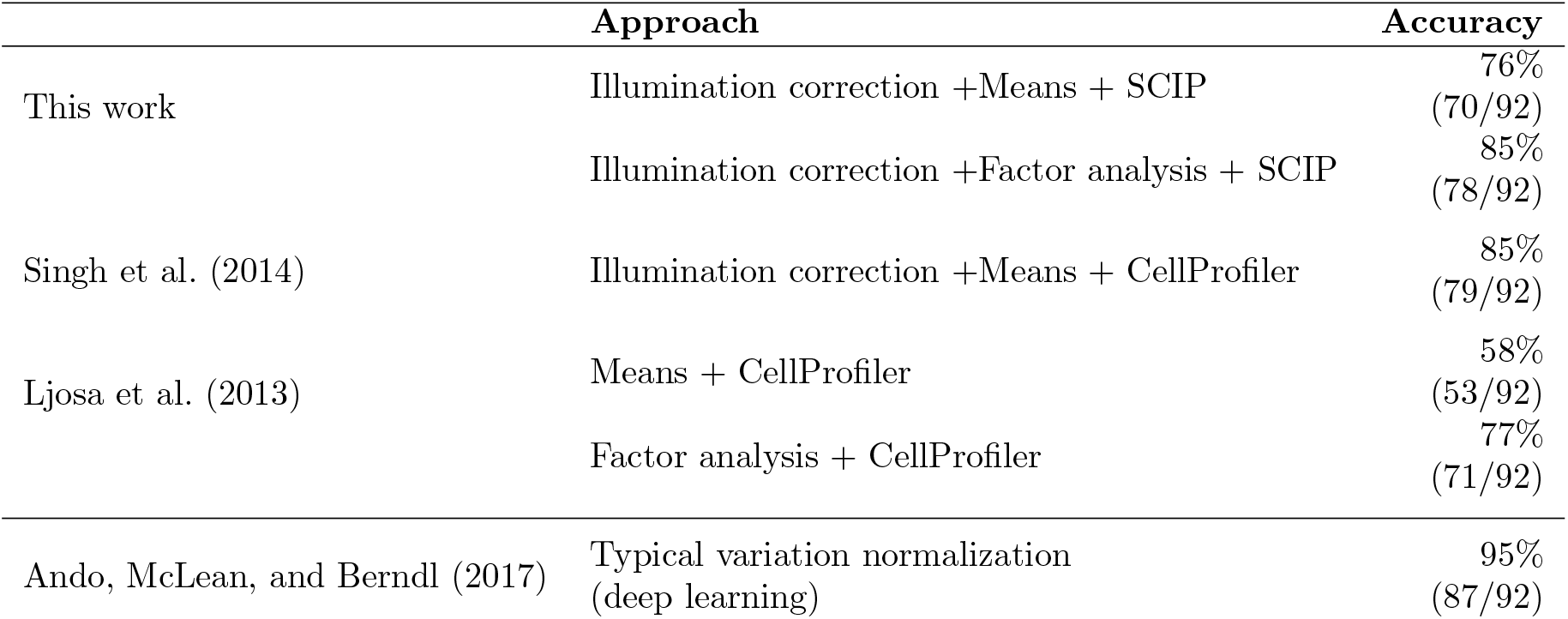
Classification accuracies obtained on the mechanism-of-action prediction task on the BBBC021 benchmark dataset using several approaches. We achieve comparable accuracies to CellProfiler-based approaches using SCIP.

We achieved a classification accuracy of 85% with features derived using SCIP following the modelling pipeline described in [6]. To validate the performance, we used the not-same-compound-or-batch cross-validation strategy proposed by Ando, McLean, and Berndl (2017). Figure 6 shows the confusion matrix. Methodological details on the classification can be found in Supplementary Section S2.1.1.

**Figure 6:**
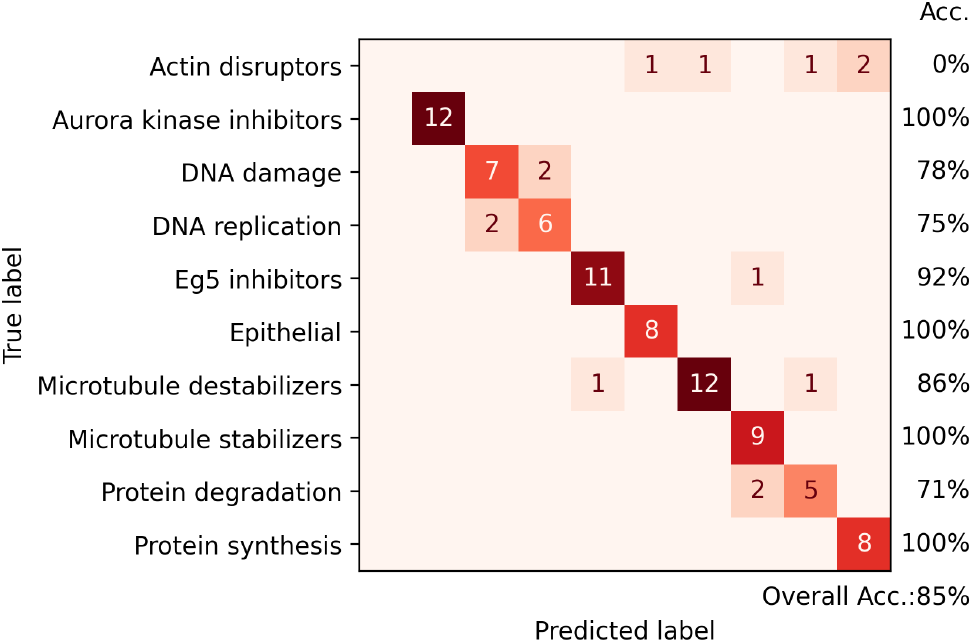
Confusion matrix for the mechanism-of-action prediction task on the BBBC021 benchmark dataset obtained with SCIP using illumination correction and factor analysis.

#### 4.1.3 Unsupervised profiling of leukocytes in human blood with automated confocal microscopy

We use SCIP to profile single cells in automated microscopy images of human whole blood cells and cluster them into 4 major cell types. We show with this use case that SCIP enables efficient profiling of cells in automated microscopy images in a distributed environment using state-of-the-art GPU-accelerated techniques.

Blood from a healthy donor was stained and imaged on a Zeiss Celldiscoverer 7 platform. The planes contain 7 channels: 4 fluorescence channels, a brightfield channel, an oblique channel and a phase-gradient contrast channel. We refer to the Supplementary Material for acquisition details. All data is accessible under study S-BIAD452 on the Bioimage Archive at https://www.ebi.ac.uk/biostudies/studies/S-BIAD505.

The images were processed with SCIP to obtain single-cell feature profiles. SCIP profiled 45,942 objects in 57 minutes and 36 seconds using 6 workers in total, of which 2 GPU-accelerated workers for CellPose-based segmentation.

We clustered cells into 4 distinct phenotypes using Leiden clustering: granulocytes, eosinophils, lymphocytes and monocytes. We also identified one unclassified cluster, which had low response for all markers in the panel. We hypothesize this cluster is made up of cell debris and platelets. Supplementary Figure S7 shows some example images of objects from the unclassified cluster. Finally, we embedded the features with UMAP to visualize the cell populations. Figure 7 shows an overview of the annotation. Supplementary section S2.1.3 describes the clustering procedure in detail.

**Figure 7:**
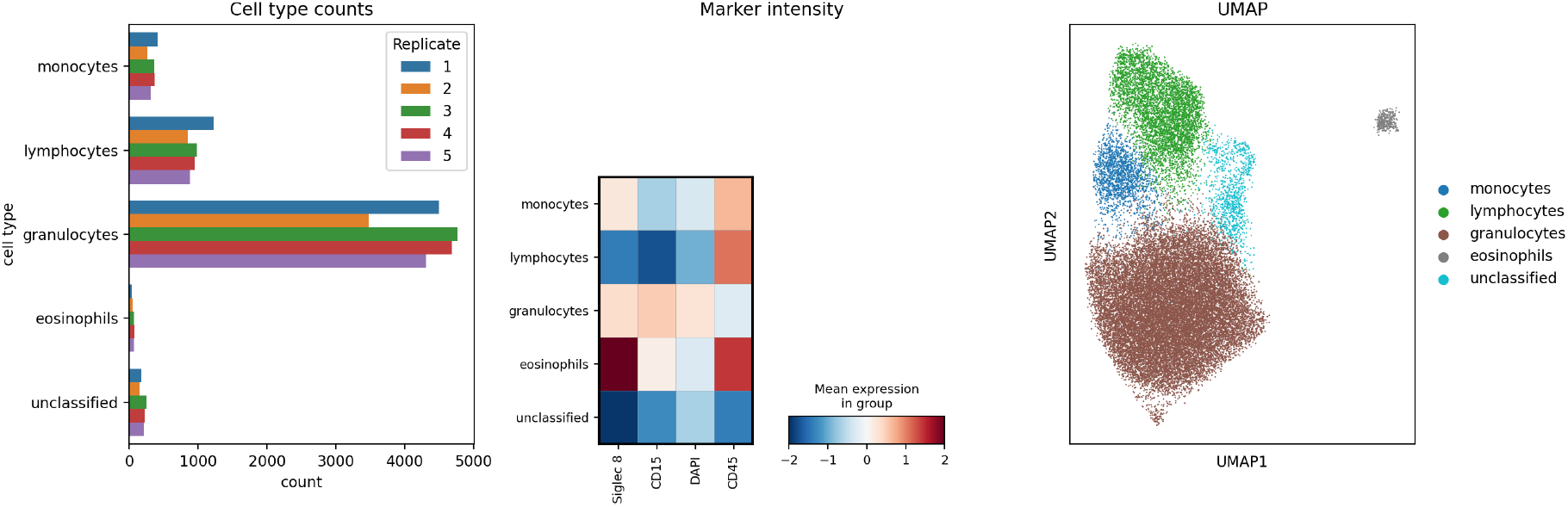
**Left** Cell type counts per replicate show an equal distribution over the replicates, as expected. **Middle** Mean expression of marker per cell type. **Right** UMAP embedding showing all populations separated in two dimensions. Note the unclassified cluster, which has low expression for all markers and is likely made up of debris and platelets.

### 4.2 SCIP scales well for increasing dataset size and number of workers

In this section, we probe the performance and scalability of SCIP in three experiments, corresponding to three parameters that determine the runtime and maximum memory usage of our approach: the dataset size *d*, the number of workers *n* and the partition size *p*. We probe the scalability of SCIP by alternatively fixing *n* or *d* and varying the other two, using the values shown in Table 5 and *p ∈* {100 · 2^*i*^ | *i ∈* {0, 1, 2, 3, 4}}. To limit the computational requirements we only used a part of the dataset from the second use case to collect the measurements. Scripts to reproduce these results can be found on Github at https://github.com/ScalableCytometryImageProcessing/benchmark.

**Table 5:** Experimental settings for number of workers and dataset size to probe the scalability of SCIP.

| Experiment | Number of workers | Dataset size | Measurement |
| --- | --- | --- | --- |
| (a) | $\{2^i \mid i \in \{1, 2, 3, 4, 5\}\}$ | 80,000 | Runtime |
| (b) | 16 | $\{1 \cdot 10^i \mid i \in \{2, 3, 4, 5, 6\}\}$ | Runtime |
| (c) |  |  | Max. memory usage |

First, we find that SCIP scales well with the number of workers. Figure 8a shows that doubling the number of workers, doubles the number of images processed per second. When the number of workers exceeds 8 the measured runtimes diverge from the ideal speedup to the incurred communication overhead between the workers. The partition size does not have a strong influence in this experiment.

**Figure 8:**
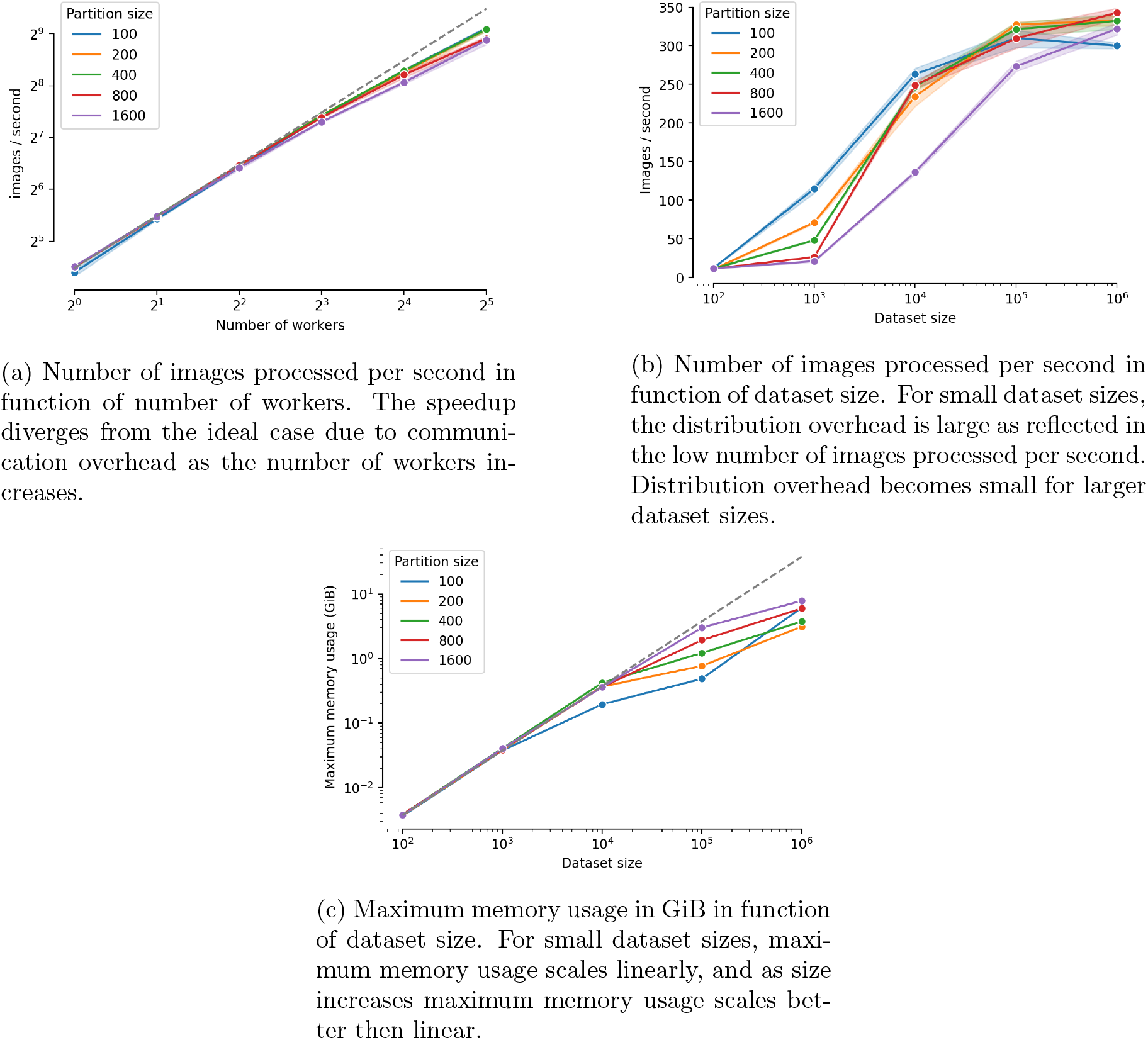
Results of scaling experiments.

Secondly, we find that SCIP scales well with respect to dataset size for a fixed number of workers. As expected, Figure 8b shows that for lower dataset sizes the distribution overhead is large, leading to a low number of images processed per second. For dataset sizes over 10 000, the overhead becomes small and the processing speed reaches a plateau of 300-350 images per second. This experiment shows an influence of partition size: for smaller datasets a smaller partition size allows for more parallelization and shorter runtimes. For larger datasets, a larger partition size is preferable to reduces communication overhead. As a rule of thumb, for datasets with less than 100 000 objects the partition size should be between 200 and 400. When going up to 1 000 000 objects, the partition size should be at least 1 000.

Finally, we find that SCIP has good scaling behaviour in terms of maximum memory usage. Figure 8c shows that memory usage scales linearly for small datasets, and sublinearly when dataset size exceeds 10 000. This is likely due to Dask’s smart task scheduling, which takes into account memory availability before scheduling new tasks, and traverses the task graph depth-first allowing for quicker reduction of the images in memory to profiles, which take up less memory. From these experiments we can conclude there is a trade-off between memory usage and speed when choosing partition size: maximum memory usage tends to be lower for smaller partition sizes, but processing speed is higher for larger partition sizes.

## 5 Availability and Future directions

SCIP is available to install from Github at https://github.com/ScalableCytometryImageProcessing/SCIP or PyPi at https://pypi.org/project/scip/. The documentation (https://scalable-cytometry-image-processing.readthedocs.io) contains usage information.

SCIP can easily be extended to be applied on other imaging data. In terms of input formats, a next step is to provide support for the standardized OME-TIFF (https://docs.openmicroscopy.org/ome-model/5.6.3/ome-tiff/specification.html) and OME-NGFF[28] file formats. This would allow users to more easily convert their data into a compatible input format to be profiled by SCIP.

An interesting use case to explore for SCIP is spatial omics data. These tissue imaging datasets are typically very large and require complex pipelines and extensive computational power to be processed. In the future, we would like to use SCIP to speed up processing of these datasets.

We would also like to extend SCIP’s profiling functionality on two fronts: First, we want to integrate CellProfiler into SCIP to make use of its extensive functionality. However, this integration is not trivial, and attempts at integration have significantly decreased SCIP’s performance. Secondly, we would like to integrate deep learning-enabled feature extraction using pretrained neural networks. These features could augment existing features for improved performance in downstream analysis tasks.

## 6 Conclusion

In this work we introduced Scalable Cytometry Image Processing (SCIP) an open-source software tool for processing large-scale image cytometry datasets implemented using the Dask framework. We discussed SCIP’s implementation and design details, highlighting the fine-grained control over task execution, scalability, and reproducibility of our software.

We demonstrated in three use cases how SCIP is used to analyze microscopy and imaging flow cytometry datasets. We reproduced earlier results obtained on stain-free classification of 8 white blood cell types, performed unsupervised clustering to profile a healthy blood sample, and reproduced results of mechanism of action-prediction using morphological features.

Finally, we show SCIP has good scaling behaviour based on timing experiments where we increased the number of workers and dataset size, as well as memory usage experiments where we measured the impact of dataset size on peak memory usage.

In conclusion, SCIP is a Python software package that enables computational biologists to extract rich feature profiles from various image cytometry datasets in a fast, reproducible and scalable manner using state-of-the-art methods.

## 7 Acknowledgments

The resources and services used in this work were provided by the VSC (Flemish Supercomputer Center), funded by the Research Foundation - Flanders (FWO) and the Flemish Government. M. Lippeveld is a Predoctoral Fellow of the Fund for Scientific Research FWO Flanders (1SB9421N). The authors want to thank Sander Thierens for his work in the initial stages of developing SCIP.

