## Supplementary Material for "A scalable, reproducible and open-source pipeline for morphologically profiling image cytometry data"

#### S1 Design and implementation

##### S1.1 Texture features

Texture features are computed using the grey-level co-occurrence matrix (GLCM) and a Sobel filter[1]. To reduce sparsity of the GLCM, $d_i$ is first binned per channel into 16 levels. From a GLCM $P$, a set of features is derived describing the texture of the image. We compute the contrast, dissimilarity, homogeneity, energy, correlation and angular second moment (ASM) as defined in Supplementary Equations S1 to S6.

$$\text{contrast} = \sum_{i,j=0}^{\text{levels }-1} P_{i,j}(i-j)^2, \tag{S1}$$

$$\text{dissimilarity} = \sum_{i,j=0}^{\text{levels-1}} P_{i,j}|i-j|, \tag{S2}$$

$$\text{homogeneity} = \sum_{i,j=0}^{\text{levels }-1} \frac{P_{i,j}}{1+(i-j)^2}, \tag{S3}$$

$$\text{ASM} = \sum_{i,j=0}^{\text{levels }-1} P_{i,j}^2, \tag{S4}$$

$$\text{energy} = \sqrt{\text{ASM}}, \tag{S5}$$

$$\text{correlation} = \sum_{i,j=0}^{\text{levels }-1} \left[ \frac{(i-\mu_i)(j-\mu_j)}{\sqrt{(\sigma_i^2)(\sigma_j^2)}} \right] \tag{S6}$$

where $\mu_i, \mu_j, \sigma_i$ and $\sigma_j$ are the means and standard deviations of $p_i$ and $p_j$.

##### S1.2 Sobel filter

The Sobel filter creates a map of $d_i$ emphasizing edges. It is computed by convolving two $3 \times 3$-kernels over over $d_i$, resulting in a vertical and horizontal approximation of the gradient, respectively $\mathbf{G}_x$ and $\mathbf{G}_y$. These are combined to compute gradient magnitude $\mathbf{G}$ as shown in

Supplementary Equation S8. We then compute the minimum, maximum, mean and standard deviation of $\mathbf{G}$.

$$\mathbf{G}_x = \begin{bmatrix} +1 & 0 & -1 \\ +2 & 0 & -2 \\ +1 & 0 & -1 \end{bmatrix} * \mathbf{d_i}, \mathbf{G}_y = \begin{bmatrix} +1 & +2 & +1 \\ 0 & 0 & 0 \\ -1 & -2 & -1 \end{bmatrix} * \mathbf{d_i} \tag{S7}$$

$$\mathbf{G} = \sqrt{\mathbf{G}_x^2 + \mathbf{G}_y^2} \tag{S8}$$

### S2 Results

#### S2.1 Use-cases

##### S2.1.1 Predicting mechanism of action for various small-molecule treatments of MCF-7 breast cancer cells

Summarizing cell profiles per treatment is either done using factor analysis (FA) or the means method, both according to [6]. In the former, an FA model is trained on a subset of the DMSO-treated cells, after which the model is used to transform the treated cells. This new representation is averaged to obtain the per-sample profile. In the latter, the profiles are immediately averaged per sample. In both cases, the median of the three replicate samples is then taken to produce a treatment profile for each compound-concentration combination.

Factor analysis describes observed variables in terms of a lower number of unobserved factors, reducing the dimensionality of the dataset. Ljosa et al. achieve optimal classification performance using 50 factors, which we also used.

##### S2.1.2 Stain-free classification of human white blood cells with imaging flow cytometry

Classification results were obtained using an eXtreme Gradient Boosting (XGB) model. We used random successive halving hyper-parameter optimization[11] validated with nested 5-fold stratified cross-validation to find optimal hyper-parameter values for the XGB model. We optimized parameters listed in Table S1. We refer to the xgboost documentation for a description of these parameters (`https://xgboost.readthedocs.io/en/stable/parameter.html`). The search was ran with 500 initial candidates and a halving factor of 2. The number of samples was increased with a factor 2 throughout the halving iterations with 5000 samples in the first iteration. Optimal configurations were selected based on the balanced accuracy.

| | **Optimal values** | | | | | **Tested** |
|---|---|---|---|---|---|---|
| | **Fold 0** | **Fold 1** | **Fold 2** | **Fold 3** | **Fold 4** | |
| **subsample** | 0.10 | 0.10 | 0.10 | 0.50 | 0.50 | 0.10, 0.20, 0.30, 0.40, 0.50, 0.60, 0.70, 0.80, 0.90, 1.00 |
| **n_estimators** | 290.00 | 260.00 | 260.00 | 270.00 | 270.00 | 10, 20, 30, 40, 50, 60, 70, 80, 90, 100, 110, 120, 130, 140, 150, 160, 170, 180, 190, 200, 210, 220, 230, 240, 250, 260, 270, 280, 290, 300 |
| **min_child_weight** | 11.00 | 19.00 | 19.00 | 19.00 | 19.00 | 1, 3, 5, 7, 9, 11, 13, 15, 17, 19, 21, 23, 25, 27, 29, 31 |
| **max_depth** | 3.00 | 4.00 | 4.00 | 6.00 | 6.00 | 6, 5, 4, 3, 2 |
| **learning_rate** | 0.10 | 0.10 | 0.10 | 0.05 | 0.05 | 0.7, 0.6, 0.5, 0.4, 0.3, 0.2, 0.1, 0.05, 0.01, 0.001 |
| **gamma** | 4.00 | 0.00 | 0.00 | 6.00 | 6.00 | 0, 2, 4, 6, 8, 10, 12, 14, 16, 18, 20, 22, 24, 26, 28, 30 |
| **colsample_bytree** | 0.50 | 0.30 | 0.30 | 0.20 | 0.20 | 0.10, 0.20, 0.30, 0.40, 0.50, 0.60, 0.70, 0.80, 0.90, 1.00 |

Table S1: Tested and optimal values used and obtained in recursive successive halving hyper-parameter optimization strategy.

To counter class imbalance, the majority class (neutrophils) was first randomly undersampled to the same level as the second most abundant class (monocytes), after which all minority classes were randomly oversampled to the same level of the monocytes.

Figures S1 and Table S2 show classification results obtained on the dataset from [24] extended with one new sample obtained using the same imaging flow cytometry protocol.

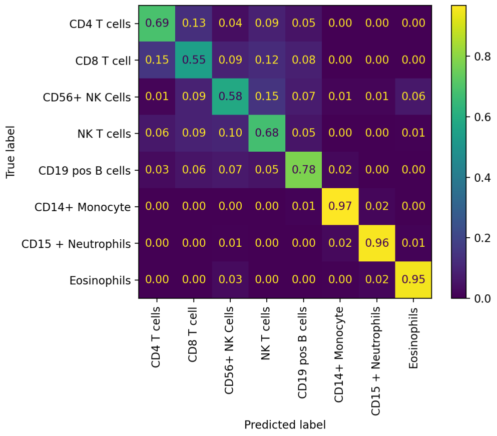


(a) Confusion matrix

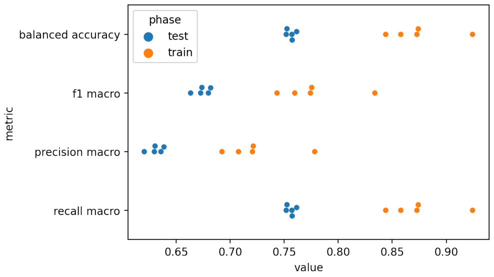


(b) Classification metrics

Figure S1: Classification results for cross-validated stain-free leukocyte classification with an eXtreme Gradient Boosting classifier.

| Phase | test | train |
|---|---|---|
| balanced accuracy | 0.769 (0.004) | 0.841 (0.012) |
| f1 macro | 0.681 (0.005) | 0.741 (0.012) |
| precision macro | 0.636 (0.004) | 0.690 (0.011) |
| recall macro | 0.769 (0.004) | 0.841 (0.012) |

Table S2: Classification metrics for cross-validated stain-free leukocyte classification with an eXtreme Gradient Boosting classifier.

###### S2.1.3 Profiling leukocytes in human blood with automated confocal microscopy

For acquisition, all cells were fixed and stained with CD45, Siglec 8 and CD15 fluorophore-conjugated antibody markers; cell nuclei were stained with DAPI. Images of cells were acquired in 5 wells, each well was imaged at 25 non-contiguous locations in 3 focal planes, each 2 micrometres apart in the Z-axis. Manual inspection flagged 12 out 125 well images with uneven illumination that could not be corrected (see Figure S2).

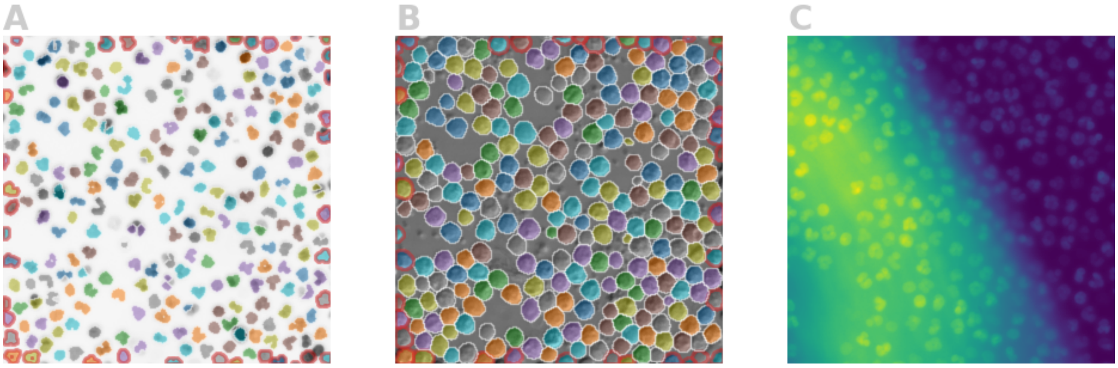


Figure S2: **A, B** Example segmentation of DAPI and oblique channel, respectively, with CellPose. Cells having a red border are discarded due to their proximity to the image border. **C** Example of image with uneven illumination, which is discarded from the dataset.

Objects whose centers were closer than 30 pixels to the border of the image were discarded. Figure S2 shows an example of a segmented image. Multiplets were gated out using aspect ratio and eccentricity as shown in Figure S3. Objects where no DAPI signal was found were also discarded. 34,838 objects remained after filtering.

Next, we aligned distributions of the fluorescence intensities across the image positions and replicates to account for technical variation. From replicate 1, 32 outliers were removed based on CD45 marker intensity and another 9 outliers based on Siglec 8 marker intensity. After this,

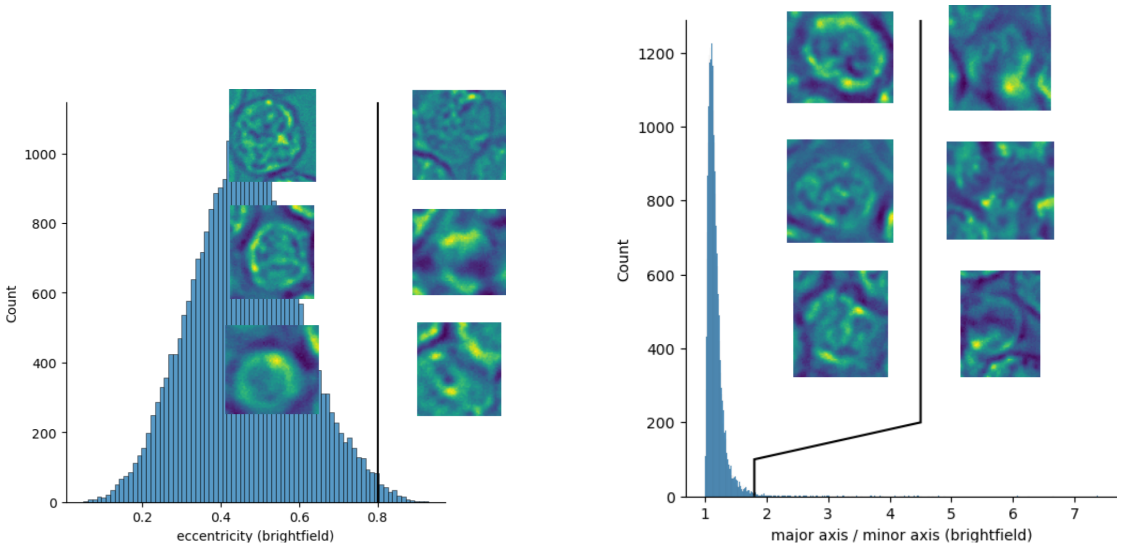


Figure S3: Multiplets were filtered out using eccentricity and aspect ratio (major axis length over minor axis length). For both figures, events with values to the right of the black line were discarded Some example images are shown as well.

an arcsinh-transform and Z-score normalization were performed per image position per replicate to align the fluorescence signals and all other intensity derived features such as signal upper quartile or mean. All other features were Z-score normalized. Supplementary Figure S4 shows the aligned intensity distributions.

Using the mutual information (MI) score, 66 features were identified and removed that had a strong relation to the replicate number after normalization. MI was computed using 30-nearest neighbors with the estimation introduced by [7].

To do so, we first filtered out multiplets and debris using the eccentricity, major axis length and minor axis length features. Features were then Z-score normalized and PCA-transformed. We constructed a 30-nearest neighbor graph based on the first 50 PCA components using UMAP distance for computing connectivity. We ran Leiden clustering with a resolution of 0.75 on this graph to discover 10 clusters. Scatter plots (see Supplementary Figures S5 and S6) of fluorescent marker intensity were used to annotate clusters with cell types.

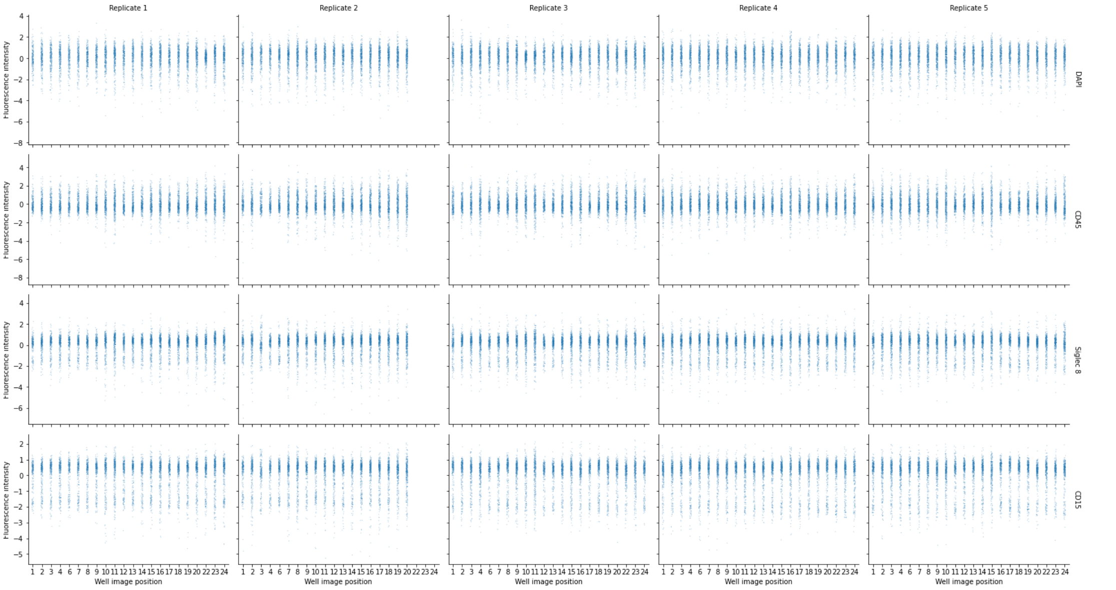


Figure S4: Strip plots of aligned intensity distributions after normalization and filtering of outlier events.

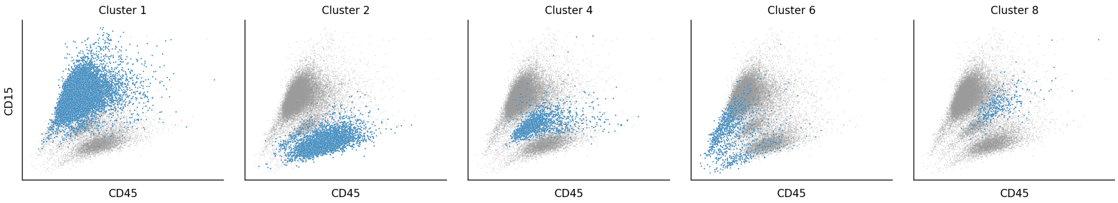


Figure S5: Scatterplots of CD45 versus CD15 intensity per cluster.

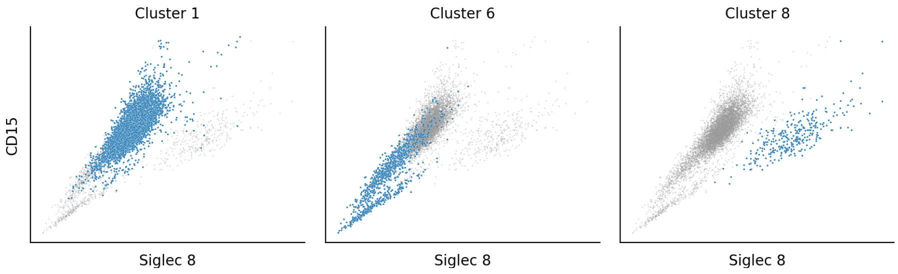


Figure S6: Scatterplots of Siglec 8 versus CD15 intensity per cluster.

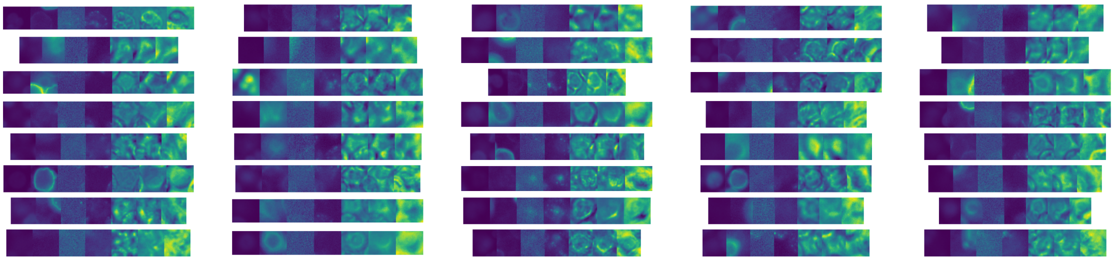

Figure S7: Example images of the unclassified cluster. Channel names from left to right are DAPI, EGFP, RPe, APC, Bright, Oblique, PGC.

| | Count | Fraction |
|---|---|---|
| granulocytes | 21725 | 0.730989 |
| lymphocytes | 4904 | 0.165007 |
| monocytes | 1737 | 0.058445 |
| unclassified | 1031 | 0.034690 |
| eosinophils | 323 | 0.010868 |

Table S3: Counts per cell type and fraction of cell type in the complete sample.
